# Behavioural state inference from movement and environmental data using Markovian step selection functions

**DOI:** 10.64898/2026.02.05.704063

**Authors:** Ilhem Bouderbala, Aurélien Nicosia, Daniel Fortin

**Affiliations:** Département de biologie, Université Laval, Québec (Qc), Canada; Mathematics Department, King Fahd University of Petroleum and Minerals, 31261, Dhahran, Saudi Arabia; Interdisciplinary Research Center for Biosystems and Machines, King Fahd University of Petroleum and Minerals, 31261, Dhahran, Saudi Arabia; Département de Mathématiques et Statistique, Québec (Qc), Canada; Centre interdisciplinaire en modélisation mathématique de l’Université Laval, Québec (Qc), Canada

**Keywords:** Animal movement, behavioural state, state-switching modelling, step selection functions, transition matrix penalty, movement ecology

## Abstract

1. Movement paths reflect temporal shifts in behavioural states, typically driven by internal and external drivers. However, the inherently multiphasic nature of these trajectories is frequently overlooked in empirical studies, an oversight that can hinder progress in our understanding of movement ecology. While Hidden Markov Models (HMMs) can successfully identify latent states—such as foraging or travelling—they face significant challenges, particularly in determining the appropriate number of states and in interpreting their ecological relevance in the context of both movement patterns and environmental covariates.
2. We present a framework based on Hidden Markov Models with Step Selection Functions (HMM-SSFs) that identifies behavioural states, represented by ecologically meaningful labels linked to explicit hypotheses about animal movement, that best explain observed movement patterns. The framework imposes interpretable conditions and diagnostic criteria on the post-identified behavioural states to ensure ecological coherence. It is grounded in the evaluation of biologically motivated scenarios rather than purely data-driven partitioning. The framework proceeds in two main steps: first, movement-based states are identified using movement-derived covariates only; second, these states are refined by incorporating environmental predictors, such as habitat structure or species interactions (e.g., predator–prey dynamics). This sequential integration enables the detection of ecological responses that are conditional on behavioural context.
3. Simulations show that the framework effectively recovers behavioural states across most conditions. State decoding accuracy was notably higher when control locations were drawn from an exponential-family distribution, compared to a uniform one. The exponential-family approach improved state separation and reduced mislabelling, especially when few control locations are generated. However, low state persistence—particularly in Encamped behaviours—resulted in an overestimation of the number of states. These findings underscore the influence of transition probabilities on behavioural labelling. Finally, we applied our framework to zebra (*Equus quagga*) movement data by combining movement predictors with changes in direction toward the nearest preferred habitat. This enabled us to distinguish between habitat-dependent and habitat-independent travelling behaviours, as well as to identify spatially finer-scale such as encamped state.
4. The proposed framework balances complexity and biological interpretability by using basic movement metrics to identify the behavioural states and their sequence that best explain multiphasic movement paths, together with environmental factors directing movement in each state. Unlike traditional methods that predefine the number of states, the framework estimates both state number and labels, offering a flexible and ecologically meaningful approach for behavioural inference.

## 1 Introduction

Movement is a fundamental process that influences not only individual fitness but also shapes population dynamics, and community structure and composition (Morales et al., 2010; Liedvogel et al., 2013). Animal movement paths reflect intricate, context-dependent responses to internal states and external environmental conditions. A wide-range of factors—including hunger, predation risk, spatial memory, locomotor capacity, and landscape structure—can influence movement behaviour (Martin et al., 2013; Bracis et al., 2015). The relative importance of these drivers often varies across space and time, resulting in movement patterns composed of multiple behavioural states expressed across diverse spatio-temporal scales (Bailey et al., 1996; Fryxell et al., 2008). For example, grazing large herbivores exhibit hierarchical foraging behaviours, moving among feeding stations within vegetation patches and across multiple feeding sites composed of such patches (Owen-Smith et al., 2010; Fortin et al., 2005b). At broader scales, their movements are typically structured around “camps”—distinct areas within their home range that encompass several feeding sites (Bailey et al., 1996).

The hierarchical nature of resource use—ranging from short-duration visits to feeding stations (<2 min) to prolonged stays at camps (1–4 weeks)—implies that movements within and between these levels should exhibit distinct behavioural signatures. Variations in basic movement metrics, such as step length and turning angle, as well as drivers of directional bias, can differentiate among behavioural states (Nicosia et al., 2017b; Prima et al., 2022). However, as movement data are collected over extended periods, both the number of meaningful behavioural states and the diversity of influencing factors tend to increase (Morales et al., 2004; Owen-Smith et al., 2010). This growing complexity poses challenges the objective identification of behavioural states and their underlying drivers, highlighting the need for analytical approaches capable of detecting states in multiphasic movement paths while maintaining both objective and biologically relevance.

Hidden Markov Models (HMMs) provide a robust framework for inferring latent behavioural states from animal movement trajectories (Morales et al., 2004; Patterson et al., 2009; Langrock et al., 2012; Michelot et al., 2016). HMMs have been integrated with both generalized biased and correlated random Walk models (BCRW) and Step Selection Functions (SSFs) to study the interplay between habitat conditions and multiphasic movement patterns. Compared to BCRW, recent studies often favour SSFs because they can be estimated with widely statistical software and provide a simple tool to evaluate multiple movement drivers simultaneously (Duchesne et al., 2015; Nicosia et al., 2017a; Klappstein et al., 2023; Hofmann et al., 2024). Despite differences, both (BCRW) and SSFs can effectively distinguish movement states and identify their ecological drivers. Notably, the two approaches have been shown to be functionally equivalent under specific control locations sampling conditions (Duchesne et al., 2015; Nicosia et al., 2017a).

HMMs are widely used to study animal movement and can be trained under various learning paradigms, depending on the availability of information about the underlying behavioural states (Wang, 2019). In a *supervised* setting, the number of states, their ecological interpretations, and the true state at each time point are all known—typically obtained through direct observation or auxiliary sensor data (Ruiz-Suarez et al., 2022). This allows direct estimation of transition and state-dependent parameters without decoding the hidden state sequence. In contrast, *unsupervised* HMMs assume no prior knowledge of the number of states, their labels, or their temporal assignments. Instead, the model jointly estimates these quantities from the data, and state labels are assigned post hoc based on ecological interpretation. In such cases, statistical criteria based on information theory or penalised likelihood methods are often used to determine the optimal number of states (Pohle et al., 2017; Dupont et al., 2025). A *semi-supervised* approach lies between these extremes: part of the state sequence is known, while the remainder is inferred (Saldanha et al., 2023). We additionally distinguish a *constrained* setting, in which the number of states and their ecological labels are specified a priori, but the exact state at each time step remains unknown and must be inferred from the data (Nicosia et al., 2017a; Prima et al., 2022; Klappstein et al., 2023). This latter case is particularly relevant when the number of states is fixed a priori to reflect ecologically meaningful behaviours (e.g., travelling vs. encamped). By contrast, fully unsupervised models are entirely data-driven, with the model structure inferred solely from the data and without explicit ecological constraints. Although such approaches can be effective for exploratory analyses, they may yield models with limited ecological interpretability. Importantly, the specification of an a priori behavioural taxonomy (e.g., “travelling” vs. “encamped”) does not guarantee that the fitted states will correspond to these labels; such correspondence must be evaluated using explicit, ecologically motivated criteria applied to the estimated state-dependent movement signatures.

In this article, we adopt an ecological guided workflow, widely used in ecological research, in which model selection is guided by predefined, biologically plausible hypotheses. Our framework relies on an *unsupervised* learning strategy with explicit ecological constraints: a limited set of candidate state numbers and their biological interpretations is defined *a priori*, while the temporal sequence of states is inferred directly from movement data. To prevent arbitrary proliferation of states, candidate models are required to satisfy predefined orderings of movement metrics, ensuring that inferred states correspond to ecologically meaningful patterns. Behavioural states are characterised using step length, angular concentration, directional persistence, and movement biases relative to habitat features, and these criteria are jointly used to infer both the number of states and their biological interpretation. Rather than seeking a unique “true” number of behavioural states, our objective is to select among a small set of ecologically motivated state taxonomies by evaluating their consistency with the data, diagnostic checks, and the expected ordering of state-dependent movement signatures. This strategy preserves ecological interpretability while avoiding both the rigidity of fully supervised models and the overfitting risks of unconstrained unsupervised approaches.

This perspective aligns with the pragmatic view advocated by Pohle et al. (2017), who showed that information criteria can be unreliable for HMM order selection and recommended combining candidate-model fitting with careful inspection of state-dependent distributions, decoded sequences, and formal model checking. Our contribution builds directly on this foundation by (i) operationalising ecologically motivated behavioural taxonomies through explicit ordering and distinctness criteria, and (ii) embedding these criteria within a reproducible, algorithmic workflow complemented by model adequacy diagnostics. While similar principles could be implemented within other directional-bias HMM formulations, we present the framework using HMMs with step selection functions (HMM-SSFs), as this formulation naturally accommodates case–control sampling, importance-sampling corrections under non-uniform availability, and direct ecological interpretation of habitat-selection coefficients.

## 2 Materials and methods

### 2.1 Hidden Markov model step selection function (HMM-SSF)

HMM-SSF characterise transitions between movement states (e.g., stationary vs. exploratory) using a hidden Markov process to capture state dependence (Nicosia et al., 2017b). For each observed step {*y*_*t*_, *y*_*t*+1_}, *J* control steps {*Z*_1,*t*_, …, *Z*_*J,t*_} are generated, where turning angles (TA) and step lengths (SL) may be sampled uniformly—i.e., *TA* ∼ 𝒰 (0, 2*π*) and *SL* ∼ 𝒰 (0, *M*), with *M* denoting the maximum inter-location distance. Alternatively, control points can be drawn from empirical or parametric distributions (Fortin et al., 2005a; Forester et al., 2009; Klappstein et al., 2023). The conditional likelihood of observing a step ending at *y*_*t*+1_, given origin *y*_*t*_ and latent state *S*_*t*+1_ = *k*, is approximated by:

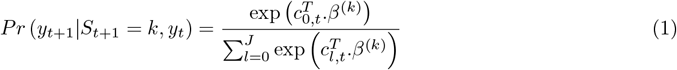

where *c*_*l,t*_ = *c* (*y*_*t*_, *Z*_*l,t*_), *Z*_0,*t*_ = *y*_*t*+1_. Here, *c* is the covariate vector comprising the SSF model covariates vector, and *β*^(*k*)^ denotes the state-specific parameters, representing the effect of each covariate on step selection. Additionally, transition probabilities may depend on *p* environmental co-variates, time of day, and interactions with neighbouring animals, modelled using a multinomial-logit link function (Morales et al., 2004; Klappstein et al., 2023).

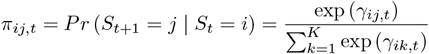

with

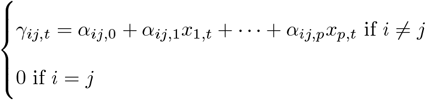

where {*x*_1_, …, *x*_*p*_} denote the covariate vector.

Klappstein et al. (2023) proposed an approach to correct for the influence of non-uniform control sampling or a low number of control points. Their method adjusts the likelihood using an *importance sampling correction*, which accounts for the bias introduced by the control distribution. In this framework, the step likelihood is approximated as:

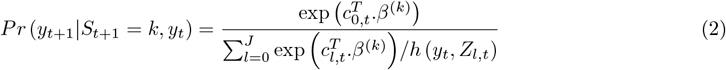

where *h*(*y*_*t*_, *Z*_*l,t*_) denotes the probability density function of the control distribution used to sample control locations. This correction ensures the likelihood properly accounts for deviations from uniform sampling, enhancing the robustness of SSF-based movement models under different control schemes. The modeller can either: (i) maximise Eq. 1 and adjust the step length and turning angle parameters, or (ii) directly maximise the corrected likelihood in Eq. 2. In the SI, we present the equivalence between the HMM-SSF model and the HMM-(biased) correlated random walk (HMM-(B)CRW.

### 2.2 Clustering metrics

Clustering metrics are used to operationalise biological hypotheses about behavioural differentiation and to guide model selection among candidate state-space models. Because behavioural modes vary across species and ecological contexts (Table 1), the choice of metrics must be tailored to the focal ecological question and the hypothesised behavioural taxonomy. For each fitted candidate model with a fixed number of states *K*, we extract a vector of clustering metrics **Ψ** from the estimated state-dependent movement parameters. These metrics typically include summaries of step lengths, turning-angle concentration, directional persistence, or context-specific measures such as movement bias toward known locations or responses to environmental risk. For basic behavioural contrasts (e.g., encamped, resting, foraging, travelling), step lengths and angular concentration are often sufficient. For example, Fryxell et al. (2008) distinguished encamped from exploratory behaviour using the frequency of directional reversals, while Ylitalo et al. (2021) identified homing behaviour through high speeds and directed movement toward a known site. More complex behavioural distinctions may require additional or context-specific metrics (Table 1; Prima et al., 2022).

**Table 1:** Examples of movement states with their description.

| Movement states | State description | Reference |
| --- | --- | --- |
| (i) Foraging<br>(ii) Searching | (i) Low angular concentration, moderate SLs<br>(ii) Concentrated TA, Relative long SLs | Fryxell et al. (2008) |
| (i) Resting<br>(ii) Foraging | (i) Shelter retreat: Low angular concentration, Short SLs<br>(ii) Food search: Relatively long SLs | Pohle et al. (2024) |
| (i) Encamped<br>(ii) Exploratory | (i) Near resource patches: Low angular concentration, Short SLs<br>(ii) Away from resources/predators: Concentrated TA, Relatively long SLs | Prima et al. (2022) |
| (i) Feeding site<br>(ii) Camp area | (i) Short SLs<br>(ii) Low angular concentration, Intermediate SLs | Bailey et al. (1996) |
| (i) Resting<br>(ii) Encamped<br>(iii) Exploratory | (i) Very short SLs<br>(ii) Low angular concentration, Moderate SLs<br>(iii) High directional persistence, Long SLs | Karelus et al. (2019) |
| (i) Resting<br>(ii) Moderate speed travel<br>(iii) Fast travel<br>(iv) Homing | (i) Low angular concentration, Short SLs<br>(ii) Intermediate angular concentration, Intermediate SLs<br>(iii) Low angular concentration, Long SLs<br>(iv) Long SLs, Movement bias towards the den | Ylitalo et al. (2021) |

Importantly, specifying an *a priori* behavioural taxonomy (e.g., “travelling” vs. “encamped”) does not guarantee that fitted states will align with those labels. Such alignment must be evaluated using explicit, ecologically motivated criteria applied to the estimated state-dependent movement signatures. In our framework, these criteria are encoded through an ordering logic defined on the clustering metrics **Ψ**. For each candidate model, we therefore evaluate whether the extracted metric vector **Ψ** satisfies the user-defined ordering logic that reflects the hypothesised behavioural structure. This logic specifies expected qualitative relationships among states—for instance, increasing step length and turning-angle concentration from resting to foraging to travelling. Model adequacy is assessed by determining whether the estimated metrics exhibit biologically interpretable separation and conform to these expectations. If the ordering logic is violated or yields ambiguous or ecologically implausible state differentiation, the candidate model is rejected and a simpler model with fewer states is fitted. This procedure is repeated iteratively until a model is obtained for which the clustering metrics satisfy the hypothesised behavioural ordering. Once a candidate model passes this validation step, final behavioural labels are assigned to states, and the most probable state sequence is decoded for the full trajectory. In this way, clustering metrics do not define behavioural states directly, but provide an explicit and ecologically grounded criterion for validating model structure and guiding state-number selection.

#### Environmental covariates for ecological state labelling

Beyond movement-derived metrics, environmental covariates play a central role in assigning ecologically meaningful labels to the inferred behavioural states. In the HMM–SSF framework, such covariates are introduced at the step-selection level and interpreted conditionally on the movement state, thereby complementing the movement-based clustering metrics Ψ.

Environmental covariates can be broadly classified into three categories. First, numerical covariates *h*(*x*) describe continuous habitat attributes measured at the step endpoint or along the step, such as elevation, vegetation density, distance to water, or predation risk. State-specific selection coefficients associated with these covariates quantify attraction or avoidance of environmental gradients and help distinguish behaviours linked to resource exploitation or risk minimisation.

Second, directional covariates capture orientation biases relative to external features through terms of the form cos(*θ* − *θ*^target^), where *θ* denotes the movement bearing and *θ*^target^ the direction toward a biotic or abiotic target (e.g., a refuge, food patch, den, or conspecific). A significant positive coefficient indicates directed movement toward the target, whereas non-significant effects suggest habitat-independent displacement. These covariates are particularly informative for discriminating habitat-dependent from habitat-independent travelling behaviours.

Third, categorical or indicator covariates encode the presence or absence of specific habitat types or landscape elements (e.g., grassland vs. woodland, inside vs. outside a protected area). In this case, state-specific coefficients measure differential use of discrete habitat classes and can be used to identify behaviours associated with habitat residency, patch use, or transitions between habitat types.

Crucially, ecological state labelling relies on the joint interpretation of movement metrics Ψ and environmental covariates, rather than on either component alone. Movement-derived quantities characterise the intrinsic locomotory mode, while environmental covariates reveal the external drivers shaping movement decisions within each state. This joint perspective ensures that behavioural states are defined not only by how animals move, but also by why and where they move, providing a coherent ecological interpretation of the inferred state structure.

### 2.3 Example taxonomy and conditions

In this section, we define the movement-based component Ψ and illustrate the proposed framework using a simple, ecologically interpretable taxonomy centred on a directional target. This choice reflects a common situation in movement ecology, where behavioural states are distinguished not only by intrinsic movement characteristics, but also by orientation toward a specific biotic or abiotic feature. We derive a set of clustering metrics Ψ:

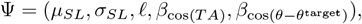

where *θ* denotes the animal’s bearing direction and *θ*^target^ represents the direction toward influential biotic or abiotic habitat attributes, such as a predator, prey, or refuge. In this formulation, *µ*_*SL*_ and *σ*_*SL*_ correspond to the mean and standard deviation of step length (SL), respectively, ℓ denotes angular concentration, *β*_cos(*TA*)_ is the coefficient associated with the cosine of the turning angle, and 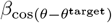 represents the coefficient associated with the cosine of the angular difference between the step direction *θ* and the direction toward the influential habitat attribute(s) *θ*^target^.

Together, these clustering metrics provide a comprehensive description of both intrinsic movement properties and environmentally driven directional responses, ensuring that the resulting states are ecologically interpretable. Step length is assumed to follow a gamma distribution, and turning angles follow a von Mises distribution—two of the most commonly used distributions in animal movement modelling (Nicosia et al., 2017b; Klappstein et al., 2023). The parameters Ψ are derived from the estimated coefficients of *SL*, log(*SL*), and cos(*TA*) obtained from the HMM–SSF model (Klappstein et al., 2023), as follows:

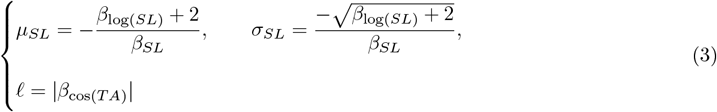

The sign and magnitude of *β*_cos(*TA*)_ provide direct insight into movement directionality: values of *β*_cos(*TA*)_ ≫ 0 indicate strong directional persistence, whereas *β*_cos(*TA*)_ ≪ 0 indicate a strong tendency to reverse direction (Klappstein et al., 2023; Hofmann et al., 2024). These clustering metrics are subsequently used to identify the states that best describe the observed movement data under the sets of conditions described below (Conditions 1–4), each corresponding to state-switching dynamics associated with a specific behavioural label (see Algorithm 1).

#### Algorithm 1

Guided model selection and state labelling for HMM-SSF

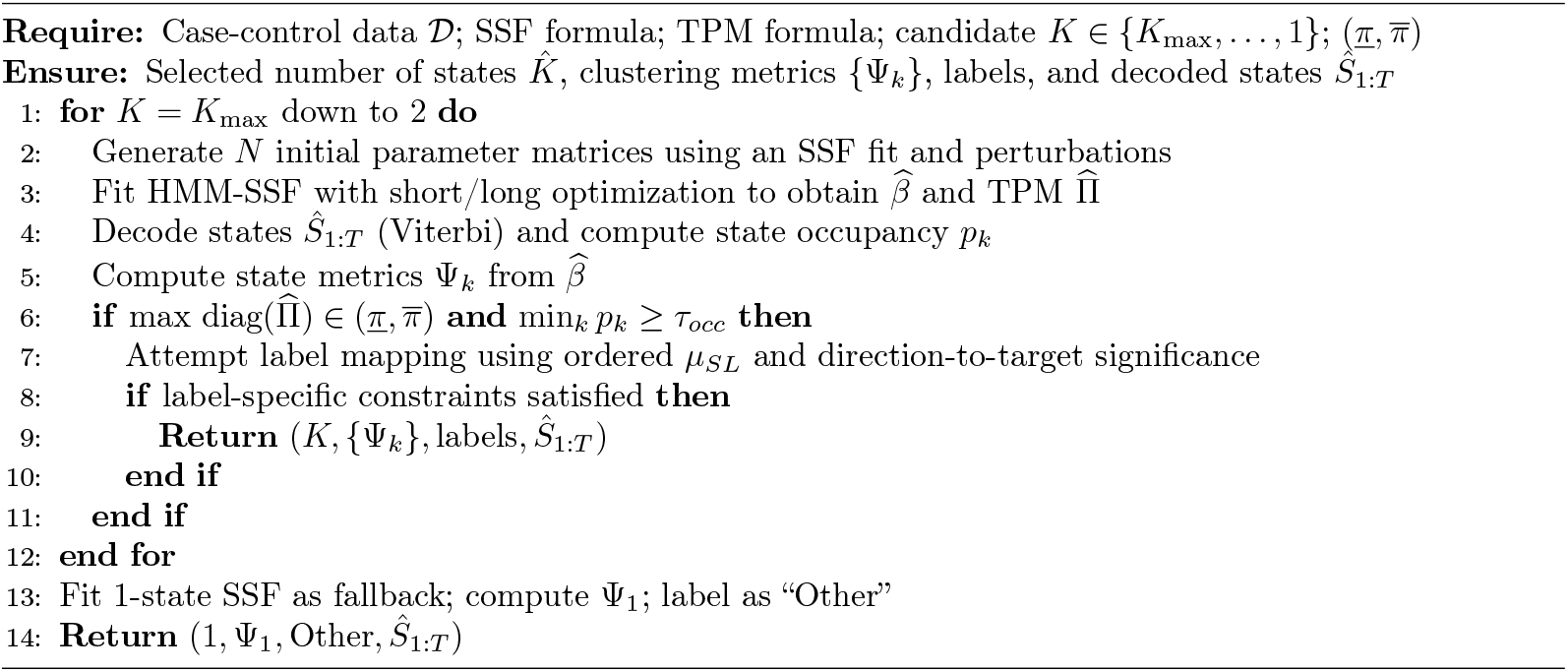

#### Four-State model (*K* = 4)

We begin by considering a model comprised of all four potential behavioural states: resting, foraging, habitat-dependent travelling (HDT), and habitat-independent travelling (HIT). The animal is assumed to remain mostly immobile when in the resting state, such that *µ*_*S*_*L* is relatively small. For simplicity, we consider that the foraging state is comprised of a single movement mode, independent of specific habitat features and without specific expectation for turning angles (*β*_cos(*TA*)_) (Fryxell et al., 2008; Nicosia et al., 2017a,b; Prima et al., 2022). However, it is worth noting that some animals employ foraging strategies—such as area-restricted search—that involve more than a single movement states, differing in movement metrics, and that can be linked to habitat structure and composition (Smith, 1974; Ward and Saltz, 1994; Fortin, 2003; Prima et al., 2022). Studies with sufficiently high-resolution movement data may find it valuable to subdivide the foraging state accordingly.

Unlike the foraging state, the travelling state is partitioned into two distinct modes. In the HDT state, movement is influenced by habitat characteristics, which may include both abiotic (e.g., topography, roads) and biotic factors (e.g., competitors, predators). In the HIT state, movement proceeds independently of the focal factors and is expected to display directional persistence (large *µ*_*SL*_ and high angular concentration *l*), such that *β*_cos(*TA*)_ *>* 0. The combination of HDT and HIT states may emerge, for example, under predator avoidance: an individual enters the HDT state when the distance to the nearest a predator is less than a given threshold (Basille et al., 2015). In such cases, the 4-state label should satisfy all conditions:

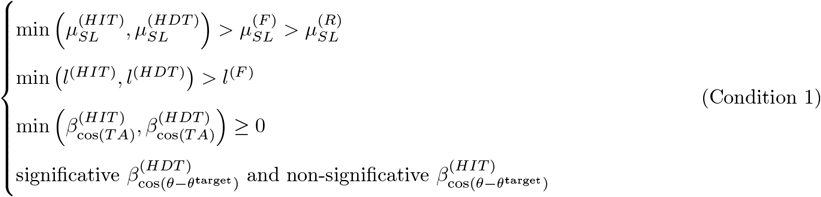

In cases where the relocation interval is short enough, the sequence of movements away from the predator should result in directional persistence also for HDT. Additionally, HIT and HDT are expected to be characterized by long step lengths. The key distinction lies in the effect of habitat on orientation: the HDT state shows a statistically significative coefficient 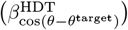, whereas the HIT state shows a near-zero, non-significative effect 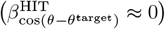. In other words, HDT reflects habitat-driven displacement—persistence combined with a directional bias toward habitat features—while HIT reflects persistence without such bias.

#### Three-state model (*K* = 3)

- **Habitat-Dependent Travelling, Foraging, and Resting**. This case applies when all inequalities in (Condition 1) are satisfied, *but* the two orientation coefficients 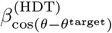 and 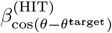 are both significative. In that situation, the two putative travelling modes cannot be distinguished—both exhibit a directional bias—so we collapse them into a single travelling state (HDT), thereby reducing the state space to three states.

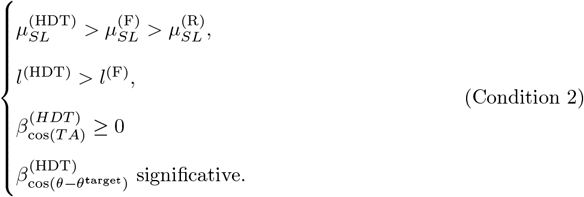

A second pathway for reducing the four–state model to three states arises when the specification is over–parametrised for the available data. We then refit the three–state model and re–assess the behavioural–state conditions. Therefore, we distinguish (i) conceptual collapsing of states when two hypothesised modes are statistically indistinguishable given the chosen metrics, from (ii) numerical or data-support limitations (non-convergence, unstable estimates), which trigger a refit with fewer states without an ecological ‘collapse’ interpretation.
- **Travelling, Foraging, and Resting**. This case applies when no environmental predictor is included to define behavioural states, or when 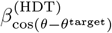 is not significative in Condition 2. The model distinguishes generic Travelling (T), Foraging (F), and Resting (R) via:

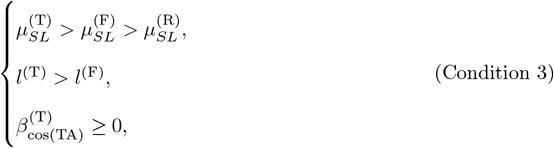

#### Two-State models (*K* = 2)

We consider in this model two states travelling and encamped. Travelling is characterised by longer SLs and higher directional persistence, whereas encamped behaviour is more variable, typically involving shorter SLs and more frequent reversals. The encamped state can encompass a mix of low-mobility behaviours and can be further subdivided into foraging and resting states (Pohle et al., 2017).

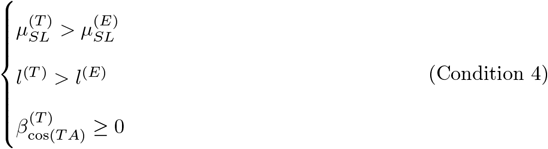

#### One-state model (*K* = 1)

For *K* = 1, the animal’s behaviour is categorised as a single, undifferentiated state designated as “Other.”

### 2.4 How to define behavioural conditions in practice

In practice, the definition of behavioural conditions should be guided by the ecological question of interest, the temporal resolution of the tracking data, the set of available covariates, and prior knowledge from the literature. The ecological question determines which behavioural contrasts are meaningful (e.g., travelling vs. encamped, habitat-dependent vs. habitat-independent movement), while the sampling interval constrains the scale at which behaviours can be resolved. For example, fine temporal resolutions may allow separation of resting and foraging behaviours, whereas coarser resolutions may only support broader distinctions such as encamped versus travelling.

The choice of covariates further informs the expected behavioural structure. Movement-only covariates (e.g., step length and turning angle) are sufficient to distinguish basic mobility modes, whereas environmental or directional covariates (e.g., orientation toward habitat features, predators, or resources) enable the formulation of hypotheses about context-dependent behaviours. These expectations should be grounded in previous empirical and theoretical studies, which provide guidance on how movement metrics and environmental responses typically differ across behavioural states (Table 1).

Based on these elements, a limited set of candidate behavioural states is defined *a priori*, together with qualitative expectations on their movement signatures—such as relative ordering of step length, angular concentration, directional persistence, or habitat-driven orientation. These expectations are not imposed as hard constraints during model fitting, but are instead used as validation criteria to assess whether the fitted state-dependent parameters are ecologically interpretable. In this way, behavioural conditions emerge from the joint consideration of ecological hypotheses, data resolution, and parameter estimates, rather than from purely data-driven clustering.

### 2.5 Decoding behavioural states in animal movement

#### 2.5.1 Pre-processing

The purpose of this first step is to prepare case-control movement data suitable for HMM-SSF analysis. This involves generating case-control data by sampling control steps from a user-specified distribution (e.g., uniform or exponential family). Covariates are also extracted to characterise both case and control locations (Fig. 1a).

**Figure 1:**
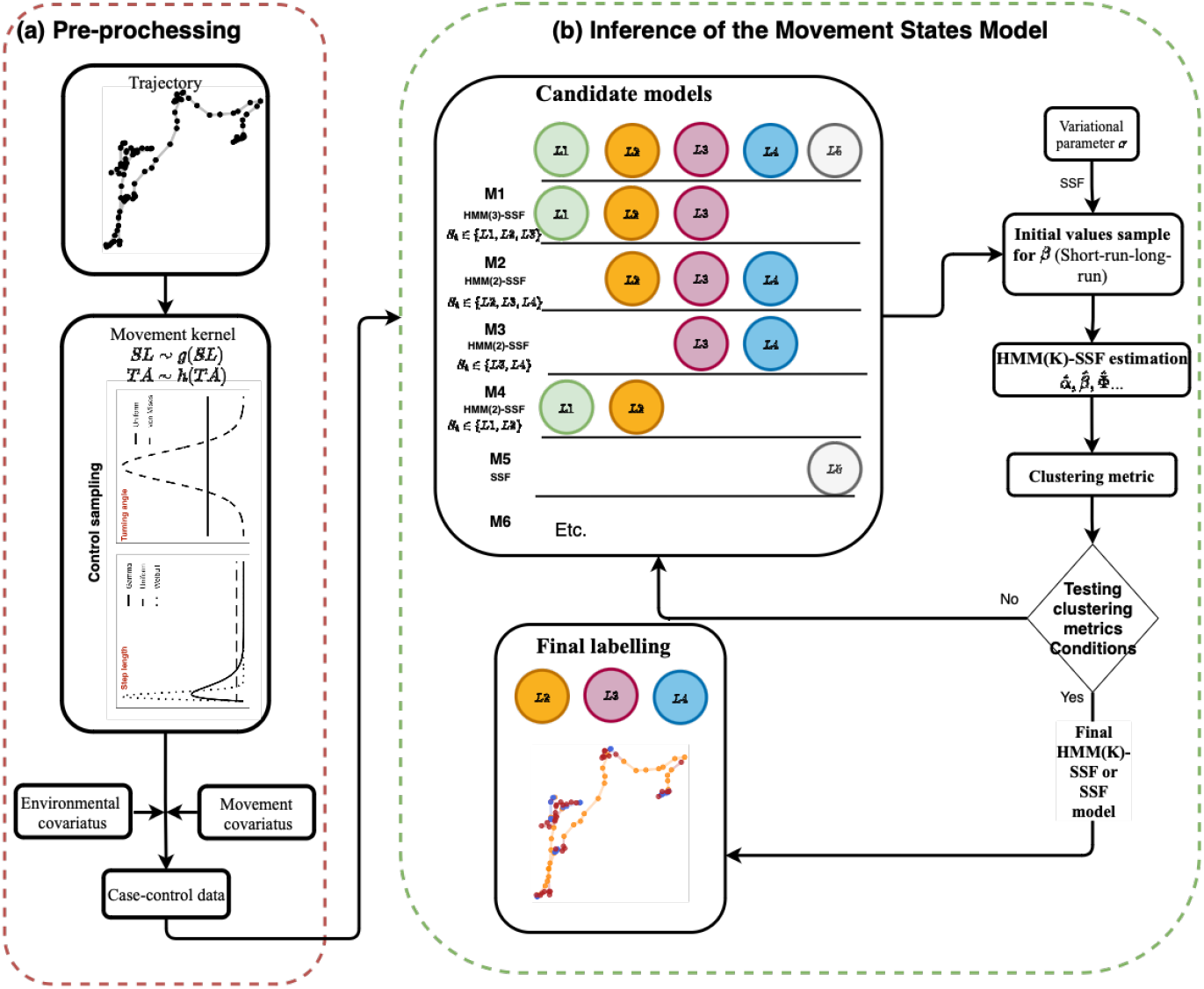
Inference diagram for estimating the number of states and assigning behavioural labels using the SSF-HMM framework. (a) Pre-processing: Preparation of case-control data, including the sampling of steps and extraction of the environmental covariates at case and control locations. (b) The movement-behaviour inference *HMM* (*K*) − *SSF* : Generation of initial values for the *β* parameters of candidate HMM-SSF models, followed by initial values sampling using the short-run/long-run procedure. The inferred states are then assigned behavioural labels based on predefined clustering metrics and an ordering logic derived from these criteria. In the provided example, five states are chosen and matched against six candidate models, with one label ultimately assigned per state.

#### 2.5.2 Short-run/long-run procedure

The aim of the second step is to generate robust initial values for the *β* parameters to explore the plausible parameter space and reduce the risk of convergence to local likelihood maxima in the HMM-SSF likelihood surface (Nicosia et al., 2017a). A single-state SSF is then fitted via conditional logistic regression, which can be implemented using the `clogit` function from the `amt` package in R (Signer et al., 2019). Based on the resulting estimates, *N* sets of *β* parameters are sampled using:

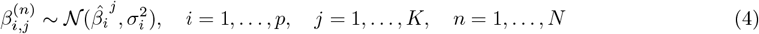

where 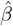 denotes the mean estimate obtained from the conditional logistic regression, and *σ*_*i*_ is a variational parameter used to simulate initial values of *β*_*i*_ around the SSF estimates. We draw *N* candidate sets of initial values from the distributions in Eq.4 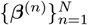 for the HMM–SSF. For each candidate, we fit the full model with a short optimisation budget (*n*_iter_short_) and record the maximised log–likelihood. We then select the best–performing run (largest log–likelihood) and restart the optimiser from its estimates using a longer number of iterations (*n*_iter_long_). This “short–run/long–run” strategy (Nicosia et al., 2017b) improves the selection of the starting values and reduces the risk of convergence to suboptimal local maxima.

#### 2.5.3 Calibration and testing

The objective of this step (Fig. 1b) is to determine the number of behavioural states and assign corresponding labels. We compile a list of candidate models that vary based on the study context corresponding to a priori hypotheses about the movement labels, and how these labels are converted into a single model.

##### Model fitting

We fit the HMM-SSF model using direct likelihood maximisation via the forward algorithm (Zucchini and MacDonald, 2009) using the initial values from the short-run/long-run procedure. The likelihood of observing a sequence of locations {*y*_1_, …, *y*_*T*_} given the model parameters is expressed as:

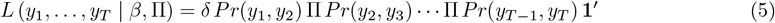

where *δ* = (*Pr* (*S*_1_ = 1), …, *Pr* (*S*_1_ = *K*)) represents the initial distribution of hidden states, *Pr* (*y*_*t*_, *y*_*t*+1_) is a diagonal matrix with the *j*^*th*^ element representing the likelihood in Eq. 2, and Π denotes the transition matrix of the Markov chain. Maximum likelihood estimates of the HMM-SSF are obtained via direct likelihood maximization using the forward algorithm, implemented using the optim optimiser in the R package hmmSSF in R (Klappstein et al., 2023). To estimate the hidden states, we can apply the Viterbi global decoding algorithm, which identifies the most probable sequence given the observations, i.e., the one that maximises *Pr* (*S*_1_, …, *S*_*T*_ |*y*_1_, …, *y*_*T*_) (Zucchini and MacDonald, 2009).

#### Testing the order in the metrics

After fitting the four–states model, if the Condition 1 are satisfied we retain the HMM(4)–SSF with *S*_*t*_ ∈ {HDT, HIT, R, F} as the final model. Otherwise, we reduce complexity using a top–down pruning scheme. We first consider a three–state HMM(3)–SSF (see Condition 2 and Condition 3), with either *S*_*t*_ ∈ {HDT, R, F} or *S*_*t*_ ∈ {T, R, F}. If the behavioural–state conditions are still not met, we fit the two–state HMM(2)–SSF with *S*_*t*_ ∈ {T, E} (Condition 4). If no HMM with *K* ≥ 2 satisfies the criteria, the model is simplified to a single–state SSF.

Candidate model space—and thus the set of behavioural labels—draws from six predefined modes {HIT, HDT, T, E, R, F}, the number of states can be defined by the user depending on the problem but we cap the number of estimated states at *K*_max_ = 4. Not all six modes can occur simultaneously: some are nested or mutually exclusive at the sampling scale (e.g., Resting vs. Encamped). Accordingly, admissible specifications are limited to biologically meaningful subsets.

### 2.6 Post-fit pruning and post-filtering rules

Our post-fit pruning and post-filtering procedure follows the same notation and logic as Algorithm 1. For each candidate number of states *K* ∈ {*K*_max_, …, 1}, we fit the HMM–SSF to obtain the SSF parameters 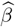 and transition probability matrix 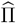, decode the most likely state sequence *Ŝ*_1:*T*_ (Viterbi), and computestate occupancy proportions 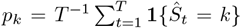. We retain the first (largest) *K* that satisfies the ensemble of constraints described below; otherwise we decrease *K* and refit.

#### 1) Numerical stability screening via TPM diagonals

To avoid degenerate transition structures, we screen fits using the diagonal of the estimated TPM. Let 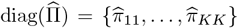. A fit is considered stable only if all self-transition probabilities lie within prescribed bounds,

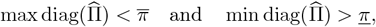

where 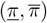 are user-specified thresholds. In our implementation, we use 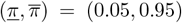. These thresholds are chosen to exclude degenerate transition regimes and can be adapted depending on temporal resolution.

#### 2) Occupancy-based pruning using Viterbi decoding

We require that the Viterbi path *Ŝ*_1:*T*_ supports all fitted states and that each state has non-negligible occupancy. Specifically, we enforce: (i) 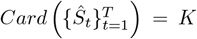 (all states are visited), and (ii) min_*k*_ *p*_*k*_ ≥ *τ*_*occ*_, where *τ*_*occ*_ denotes the minimum occupancy threshold. We set *τ*_*occ*_ = 0.05. If either condition fails, the model is refitted with *K* ← *K* − 1.

#### 3) Interpretability: ordering, distinctness, and label-specific constraints

For each candidate *K* that passes the TPM and occupancy screens, we compute state-level metrics {Ψ_*k*_} from 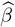 (as in Algorithm 1) and attempt a label mapping based on movement summaries derived from 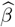 (e.g., ordered mean step length *µ*_*SL*_ and directional persistence proxies). The mapping is accepted only if the *label-specific constraints* encoded are satisfied, namely:

- **Ordering constraints:** labels are assigned consistently along the ordered *µ*_*SL*_ gradient (larger *µ*_*SL*_ corresponding to faster displacement modes, smaller *µ*_*SL*_ to slower/resting modes).
- **Distinctness constraints:** candidate labels must correspond to states that differ meaningfully in their estimated movement/SSF signatures (e.g., contrast in *µ*_*SL*_ and in key directional coefficients used by the code to separate targeted vs. non-targeted movement and foraging/resting-type states).

If the label mapping fails these constraints for a given *K*, we decrease *K* and repeat the fitting and filtering.

#### 4) Final selection and fallback

The selected number of states 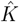 is the first (largest) *K* encountered in the descending search that satisfies the stability condition on 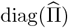, the occupancy constraint min_*k*_ *p*_*k*_ ≥ *τ*_*occ*_, and the label-specific constraints. If no multi-state solution satisfies these requirements, we fit a 1-state SSF as fallback, compute Ψ_1_, and assign the label “Other”, returning (1, Ψ_1_, Other, *Ŝ*_1:*T*_).

##### User responsibility

Because behavioural labels remain scientific constructs, users must verify that the final output (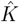, {Ψ_*k*_}, labels, *Ŝ*_1:*T*_) is biologically coherent for the study system. In practice, we recommend inspecting the retained 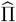, the occupancy proportions {*p*_*k*_}, and the state-specific movement summaries implied by 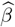 (e.g., step-length and turning-angle patterns) to confirm that the selected labelling is consistent with domain knowledge.

### 2.7 Simulation study

Following the principles outlined in Section 2.3, we simulated the movement of computer agents across five scenarios (Table. 2). Each scenario was designed to evaluate the framework’s ability to: (1) detect the true number of behavioural states, (2) accurately assign behavioural labels to those states, and (3) reliably reconstruct the sequence of behavioural states along movement trajectories over time. We also assessed the framework’s sensitivity to variations in state transition probabilities and the choice of sampling distribution for generating control locations based on the following distributions:

**Table 2:** Parameter values {*µ*_*SL*_, *σ*_*SL*_, *κ, l*, Π} used to simulate movement paths across five scenarios, with varying numbers of states (*K*) and the transition probabilities.

| Scenario | Labelling | State | $\mu_{SL}$ | $\sigma_{SL}$ | $\kappa$ | $l$ | $K$ | $\Pi$ |
| --- | --- | --- | --- | --- | --- | --- | --- | --- |
| 1 | Other (Travelling) | O | 15 | 5 | 10 | 10 | 1 | – |
| 2 | Travelling, Encamped | T<br>E | 15<br>3 | 5<br>1 | 10<br>3 | 10<br>3 | 2 | $\begin{pmatrix} 0.8 & 0.2 \\ 0.4 & 0.6 \end{pmatrix}$ |
| 3 | Travelling, Encamped | T<br>E | 15<br>3 | 5<br>1 | 10<br>3 | 10<br>3 | 2 | $\begin{pmatrix} 0.2 & 0.8 \\ 0.4 & 0.6 \end{pmatrix}$ |
| 4 | Travelling, Encamped | T<br>E | 15<br>3 | 5<br>1 | 10<br>3 | 10<br>3 | 2 | $\begin{pmatrix} 0.8 & 0.2 \\ 0.8 & 0.2 \end{pmatrix}$ |
| 5 | Travelling, Resting, Foraging | T<br>R<br>F | 15<br>0.3<br>5 | 5<br>0.1<br>3 | 10<br>0.5<br>-5 | 10<br>0.5<br>5 | 3 | $\begin{pmatrix} 0.8 & 0.15 & 0.05 \\ 0.1 & 0.7 & 0.2 \\ 0.2 & 0.1 & 0.7 \end{pmatrix}$ |

1. Case (GU_vMU): This corresponds to uniform sampling for controls. Movement is modelled using a gamma distribution for SL and a von Mises distribution for TA.
2. Case (GG_vMvM): In this case, both movement and control distributions are modelled using gamma-distributed SLs and von Mises-distributed TAs.

For the uniform sampling, we compared the results for *J* = 20 and *J* = 200 to examine the effect of the weak law of large numbers on the convergence from HMM-SSF to HMM with GRW (Nicosia et al., 2017a).

For each scenario, movement paths were generated using a Hidden Markov Model incorporating a correlated random walk, with

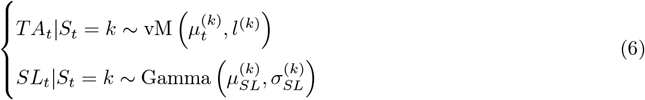

Here, 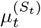 and 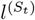 denote the direction and angular concentration of the vector

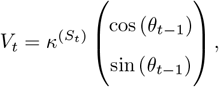

here, *θ*_*t*−1_ denoted directional persistence from the previous step, so in context of CRW 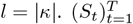 first generated from a discrete-time first-order Markov chain with a predefined transition probability matrix **Π** and an initial state distribution *δ*. For each simulated trajectory, the initial state *S*_1_ was drawn from uniform distribution. The value of the transition matrix under each scenario is defined in Table. 2

In the simulation study, we applied Condition 3 to Scenario 5 (Travelling–Resting–Foraging) and Condition 4 to Scenarios 2–4 (Travelling–Encamped).

## 3 Results

### 3.1 Framework verification: simulation study

#### 3.1.1 Number of states and control sampling distribution

The simulated movement trajectories (Fig. 2) exhibited the expected behavioural patterns: scenarios involving Travelling show larger ranges and longer step lengths (SLs), whereas distinct turning angles (TAs) differentiate Travelling from Foraging behaviours.

**Figure 2:**
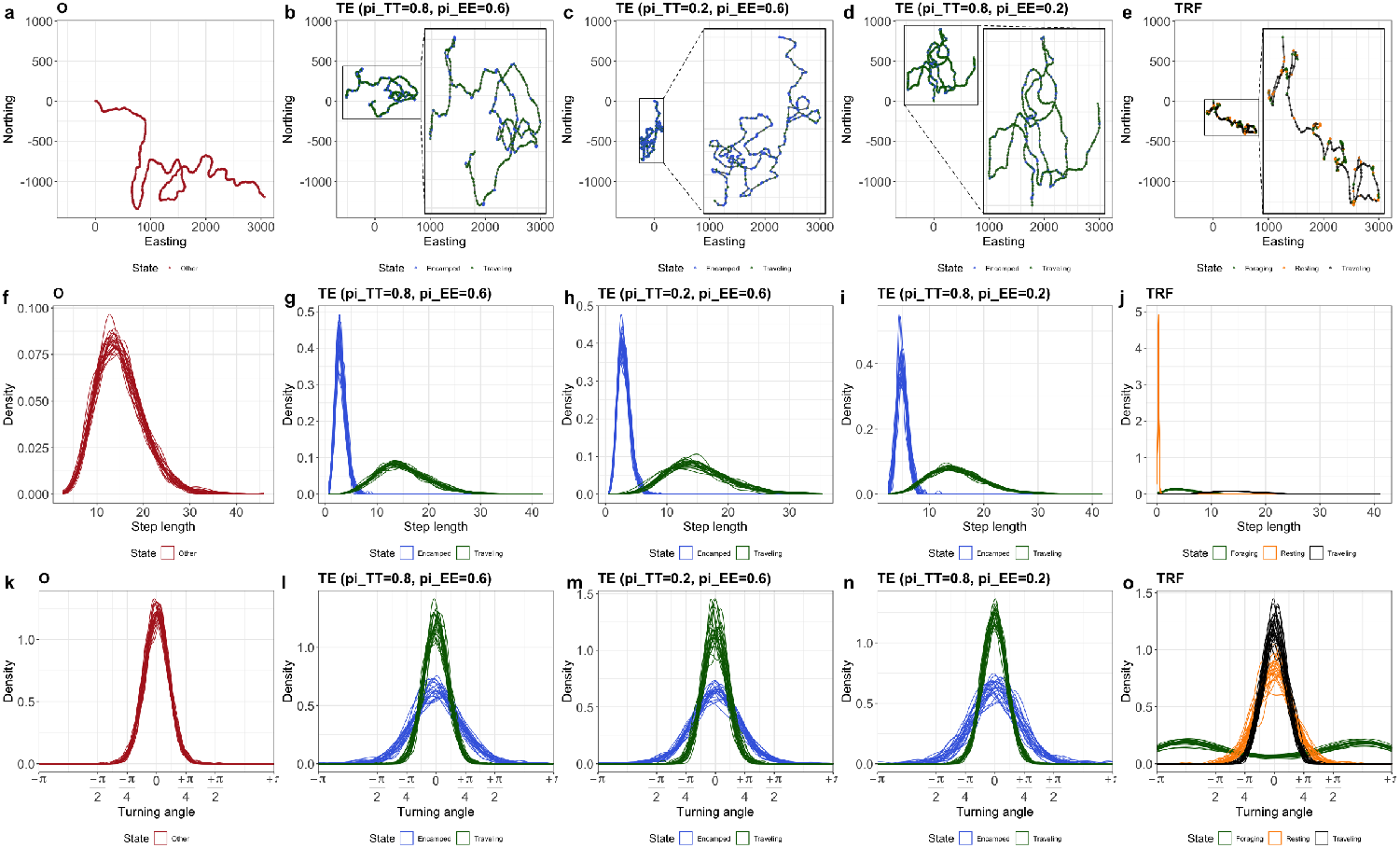
Simulation results for five scenarios. Columns represent: Scenario 1 (O), Scenario 2,3 and 4 (three variants with different *π*_*TT*_, *π*_*EE*_), and Scenario 5. Rows display: (1) a single simulated trajectory,(2) SL distributions across 25 individuals, and (3) TA distributions across 25 individuals. Colours indicate behavioural states. Each simulation ran for 500 steps.

Across scenarios, the framework reliably recovered the correct number of behavioural states (one to three) for almost all individuals; the only consistent deviation occurred under the GU_vMU sampling scheme (*J* = 20). Only one individual under the GG_vMvM sampling and one individual under the GU_vMU (*J* = 200) sampling were misclassified as belonging to a three-state model within Scenario 4 (TE with *π*_*TT*_ = 0.8 and *π*_*EE*_ = 0.2). Additionally, six individuals were misclassified under the GU_vMU (*J* = 200) sampling method with the three-state model (scenario TRF) and considered belonging to two-states model. Uniform sampling with *J* = 20 performed the worst; for example, in the three-state scenario, only 8% were correctly classified, compared to 100% with gamma and von Mises sampling (Table. 3). Similar trends occurred in two-state scenarios. The case of gamma and von Mises (GG_vMvM) with *J* = 20 was almost equivalent to uniform sampling but with a large number of control points but with better performance.

**Table 3:**
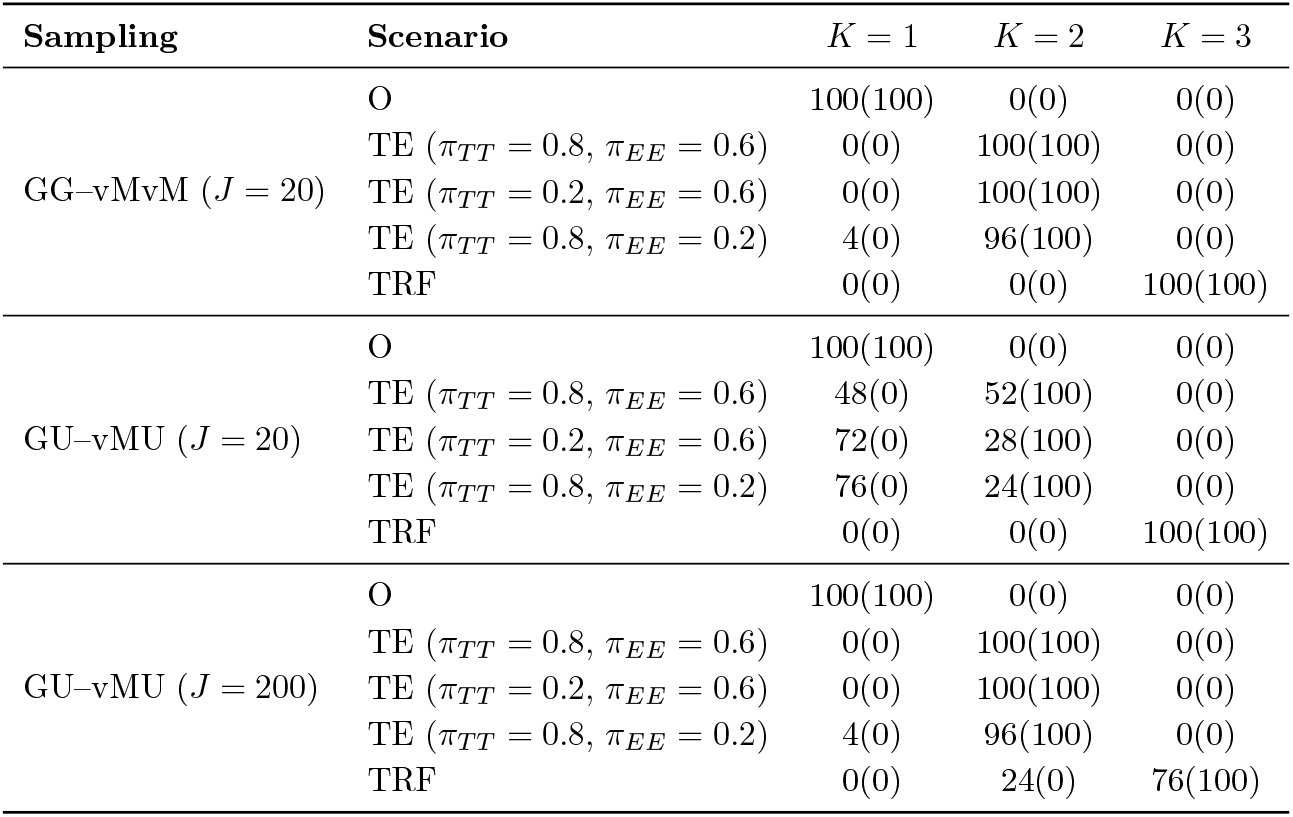
The percentage of individuals with correct matching between the estimated number of states and ground truth under the six scenarios: Scenario 1 (Other), Scenario 2, 3 and 4 for *π*_*TT*_ = 0.8, *π*_*EE*_ = 0.6, *π*_*TT*_ = 0.2, *π*_*EE*_ = 0.6, and *π*_*TT*_ = 0.8, *π*_*EE*_ = 0.2, and Scenario 5 (Travelling-Resting-Foraging). Results are shown for three choices of the SL and TA distributions for the controls: GG–vMvM, GU–vMU with *J* = 20, and GU–vMU with *J* = 200. Values in parentheses indicate the true percentages.

#### 3.1.2 State labelling and control distribution

The identification of the correct state label (Figs. 3) was least successful with uniform control point sampling with *J* = 20. For instance, in Scenario 3, the model incorrectly identified 72% and 73% of observations corresponding to encamped and travelling as one-state model (considered as a travelling mode). Similarly, in Scenario 2, encamped state labelling was only 19% accurate with uniform sampling at *J* = 20, compared to 99% accuracy with gamma and von Mises distributions (GG_vMvM (J=20)), as well as uniform sampling with *J* = 200 (GU_vMU (J=200)). However, the performance of gamma and von Mises distributions for SL and TA remained highly accurate across all scenarios, with a maximum correct identification rate of 100% and a minimum of 98% for the foraging state in Scenario 6 (travelling-resting-foraging, TRF) (Figs. 3). With uniform sampling and *J* = 200 control points, performance under the three-state (TRF) model is slightly lower: the correct identification rates are 69% for foraging and 66% for resting.

**Figure 3:**
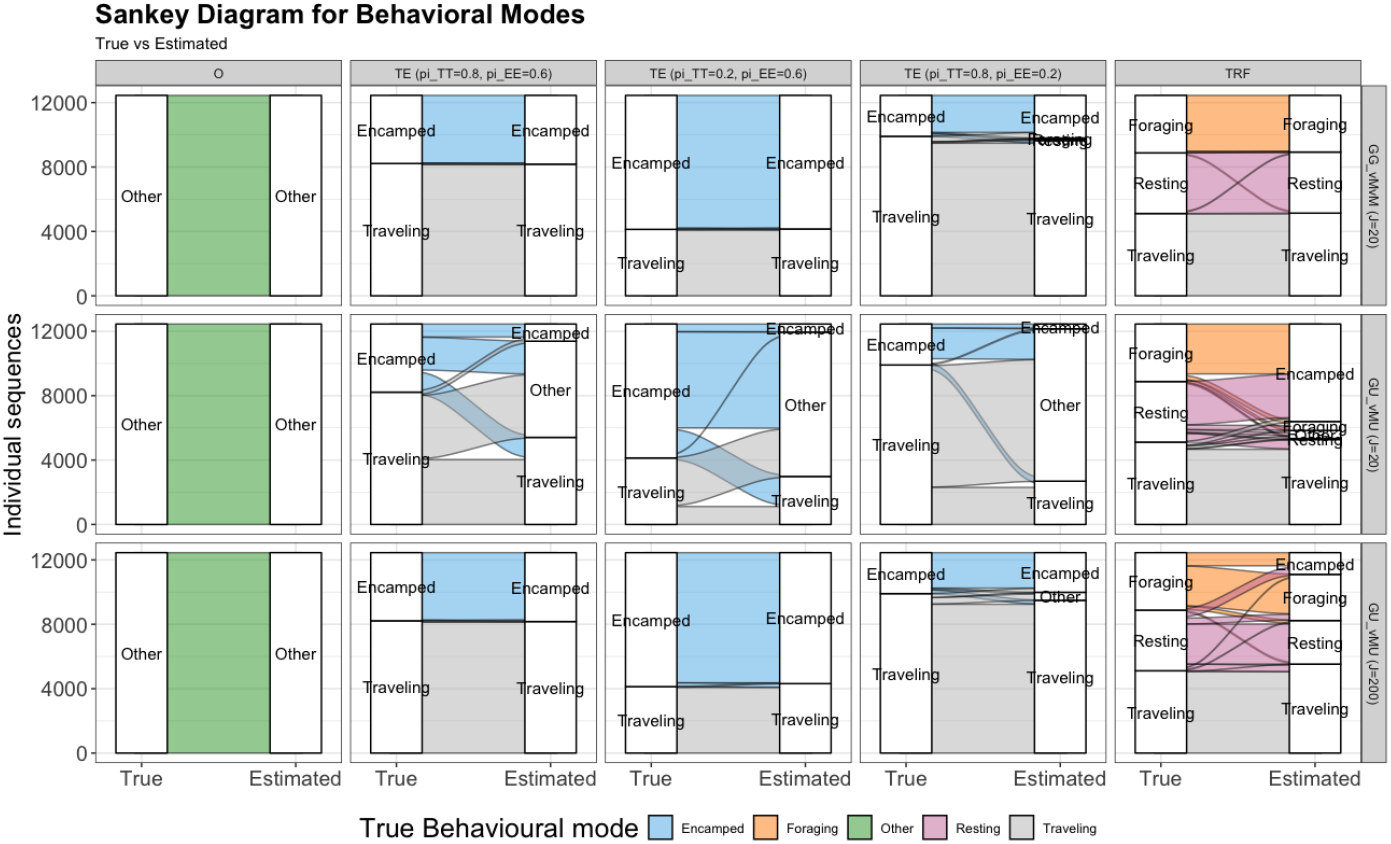
Sankey diagrams comparing **true** and **estimated** (using viterbi decoding algorithm) behavioural modes across simulation scenarios and data regimes. Columns correspond to scenarios (O; TE with transition probabilities (*π*_*TT*_, *π*_*EE*_) ∈ {(0.8, 0.6), (0.2, 0.6), (0.8, 0.2)}; TRF), and rows to data regimes (GG_vMvM *J* = 20, GU_vMU *J* =20, GU_vMU *J* =200). In each panel, flows connect true labels (left) to model-estimated labels (right); band width is proportional to the number of individual sequences. Colours denote modes: Encamped (blue), Foraging (orange), Other (green), Resting (pink), and Traveling (grey). The diagrams visualize both correct assignments (dominant straight flows) and confusions between modes (crossing flows), highlighting how mislabelling patterns vary with Π and sample size *J*.

We observed that Scenario 4, with transition probabilities *π*_*TT*_ = 0.8 and *π*_*EE*_ = 0.2, was the least accurate under both GG_vMvM (J=20) and GU_vMU (J=200) control distributions. In this scenario, the travelling state was correctly identified 93% of the time, and the encamped state was correctly identified 87% of the time under the GU_vMU (J=200) distribution. The lower accuracy in this case is mainly due to the misspecification of the HMM(2)-SSF model as an SSF model or as a two states model but incorrect labelling, leading to 4% of encamped states being misclassified as “other” state and 10% as travelling state (Figs. 3 and Table. 3). The Sankey diagrams in Fig. 3 compare true versus estimated behavioural modes. Under the Gamma–von Mises control distributions, assignments are nearly perfect with only minor mislabeling. A similar pattern holds for uniform sampling with *J* = 200, except in the three-state (TRF) scenario, where we observe noticeable mixing—primarily foraging as encamped and resting as encamped, foraging, and travelling. In this case, the correct identification rates drop to 69% for foraging and 66% for resting.

#### 3.1.3 State labelling and transition probability

Reducing the probability of staying in the travelling state (*π*_*TT*_) from 0.8 to 0.2 (Fig. 3: Scenarios 2 and 3) had minimal effect on travelling state labelling under both GG_vMvM (J=20) and GU_vMU (*J* = 200) control distributions. However, within the GU_vMU (*J* = 200) distribution, it led to a slight decrease in the percentage of correctly identified encamped states from from 99% to 97%.

Reducing the probability of remaining in the encamped state (*π*_*EE*_) from 0.6 to 0.2 (Scenarios 3 and 4) had a more pronounced impact on labelling accuracy for both states. Specifically, under the GU_vMU (*J* = 200) control point distribution, the accuracy of travelling state labelling dropped from 99% to 93%, while encamped state labelling accuracy decreased to 87%. This decline in accuracy is partly attributable to model misspecification, but also to the mislabelling of 10% of encamped points as travelling states in Scenario 4. This suggests that transitions from the encamped to the travelling have a stronger influence on labelling accuracy than transitions in the opposite direction.

### 3.2 Framework application: case study

Here, we apply the framework presented in section 2.3 to an existing dataset consisting of GPS locations of plains zebras (*Equus quagga*) collected every 30 minutes between January and April 2014 (n = 7,246 locations from one individual) in Hwange National Park, Zimbabwe Michelot et al. (2019); Klappstein et al. (2023). We added a habitat covariate, defined as the cosine of the difference between the animal’s movement direction and the direction toward the nearest grassland habitat. This covariate was included to test whether the two travelling states, HDT and HIT, could be distinguished based on their significance, as well as to evaluate the sign of 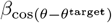.

We define the log-linear component of the state-specific likelihood in Eq. 1 by

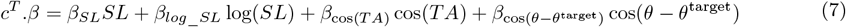

For the parameters used in the short-run/long-run procedure (Section 2.5.2), we considered a number of replications *N* = 10. The variation scale parameter (*σ*) was set to 1 for all parameters except for step length (SL), where *σ* was set to 4. The number of iterations for the short run was set to *n*_iter_short_ = 100, and for the long run, it was set to *n*_iter_long_ = 1000.

The framework identified three behavioural states: Habitat-Dependent-Travelling (HDT), Foraging and Resting states. These states were well discriminated, as evidenced by smoothed state probabilities (Fig. 4a) and the Viterbi-decoded state sequence (Fig. 4b). The HDT state (blue line) represents directed movement toward the grassland habitat 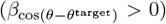, whereas the Foraging and Resting states (green and red lines, Fig. 4b) reflect localised behaviour. The HDT and Foraging and Resting states strongly differ in SL and TA distributions (Fig. 4c and 4d):

**Figure 4:**
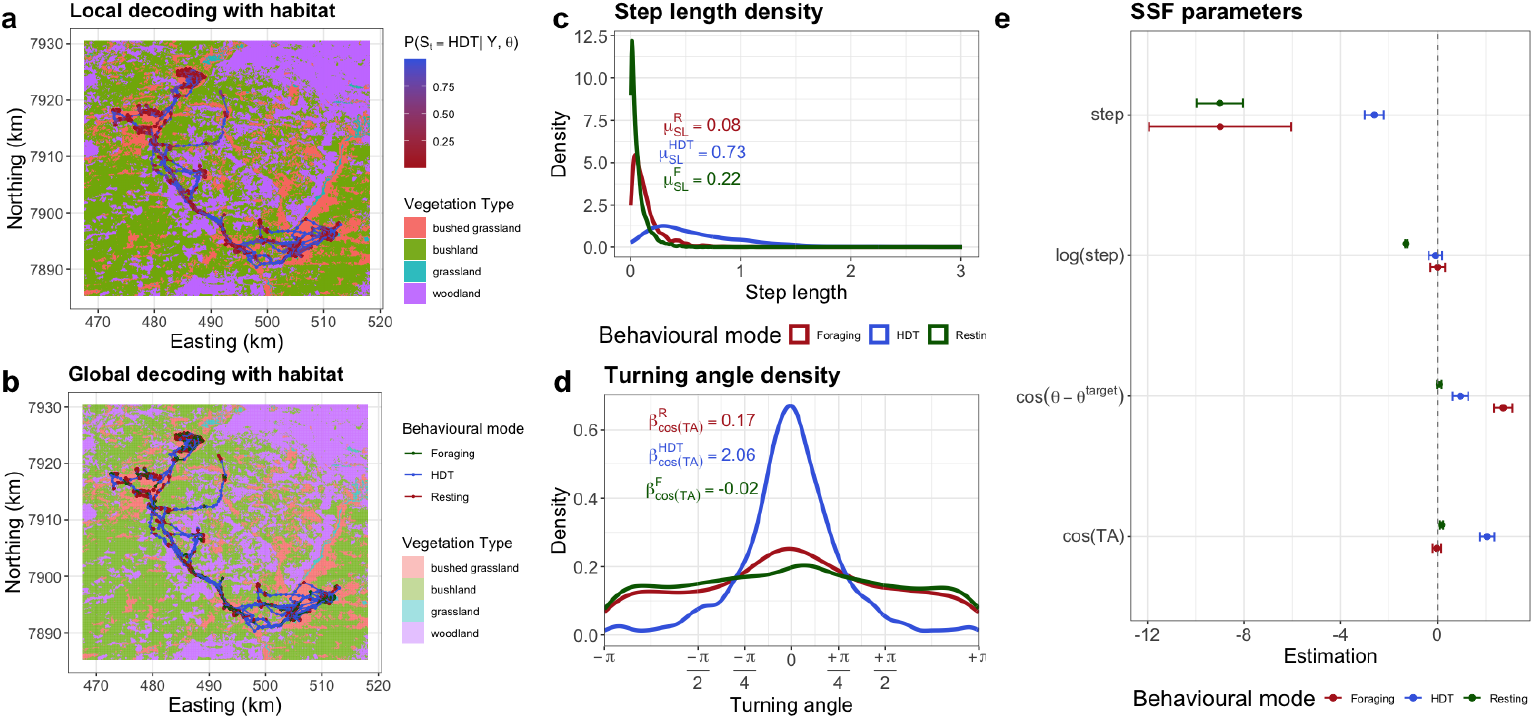
The framework applied to zebra geospatial data. **a** Smoothed probability of being in the HDT mode, *Pr* (*S*_*t*_ = *HDT* |*y, β, α*, Π), contrasted with vegetation types. **b** Viterbi global decoding. **c** SL density for the three states. **d** TA density for the three states. The three densities were estimated based on the the Viterbi decoding. **f** The confidence intervals for vegetation type (see S2 for estimation details).

1. **HDT:** 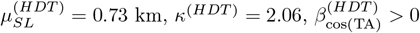 and significative 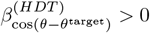.
2. **Foraging:** 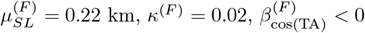 and significative 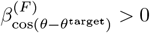
3. **Resting:** 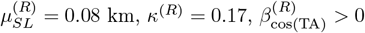 and 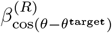 not significative.

We observed that the HDT state was meaningful; thus, in the three-state model (Condition 2), 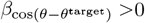 for both HDT and Foraging states. This indicates that animals were attracted toward grassland habitat in both modes. However, the parameter was not significative in the Resting state, consistent with the fact that animals are not actively searching for resources when resting.

The results differed from those reported in Klappstein et al. (2023), where the two states considered were Travelling and Encamped. This discrepancy can be explained, first, by differences in the environmental variables included in the SSF: in our case, we incorporated the direction toward the preferred grassland habitat of zebra. Moreover, unlike their approach, which predefined the number of states, our method estimated both the number of states and their labels simultaneously. In Klappstein et al. (2023), they also proposed a three-state model; our analysis empirically supports this three-state structure, as just demonstrated.

## 4 Discussion

Characterising behavioural states from animal movement data remains a central challenge in movement ecology, particularly in the absence of direct individual-level observations. We introduce a framework that integrates HMMs with step selection functions (SSFs) to infer both the number and interpretation of behavioural states characterizing the multiphasic structure of movement paths. The framework describes the order of selection in an unsupervised learning with ecological constraints by leveraging ecologically interpretable movement metrics—such as step length, angular concentration, and directional persistence—as clustering criteria. This design enables biologically grounded especially when the number or nature of states is uncertain. To balance model complexity and interpretability, the number of states is limited (Schmidt et al., 2016), as the number of possible models grows exponentially with each additional state (e.g., four models for three states, fourteen for four states; see S1).

Our framework also addresses practical challenges in applying SSFs to movement data, particularly regarding control point generation—a key factor that can bias parameter estimates if improperly selected (Forester et al., 2009; Fieberg et al., 2021). We compared two sampling strategies: uniform and exponential family distributions. Uniform sampling with a small number of control points (e.g., *J* = 20) often yields poor state differentiation, whereas increasing *J* to 200 or using parametric distributions (e.g., gamma for step length, von Mises for turning angle) substantially improves inference. This improvement reflects convergence to GRW behaviour as *J* → ∞ (Duchesne et al., 2015; Nicosia et al., 2017a). Given the computational cost of large *J*, we recommend parametric sampling to achieve comparable accuracy with fewer control points. In addition, the choice of a small number of control points is related to examining the equivalence between the HMM-SSF and the GRW counterpart. if equivalence with GRW is not of interest, one may choose J based on accuracy-cost trade-off, not on the asymptotic equivalence argument. Simulations showed that low persistence in certain states—such as encamped behaviour—can result in underestimation or mislabelling of those states. This supports the use of semi-Markovian transition structures to enhance model realism by better representing state duration, as proposed in earlier studies (Langrock et al., 2012; van de Kerk et al., 2015).

In this study we specify a small set of candidate state taxonomies (e.g., Travelling–Encamped; Travelling–Resting–Foraging; and, habitat-dependent vs. habitat-independent travelling) and pair them with state-specific movement signatures and environmental drivers. Density-independent cues (orientation to landscape features and habitat structure) anchor displacement behaviours and cleanly separate habitat-dependent from habitat-independent travelling. Compared with information-criterion workflows that optimise fit but can yield behaviourally opaque states (Pohle et al., 2017), our constrained, unsupervised strategy prioritises biologically defensible partitions while still estimating the latent sequence. This complements pragmatic order-selection tools by offering an ecologically anchored alternative when “best AIC/BIC” and “best biology” diverge (Link and Barker, 2006; Aho et al., 2014). It also accords with model-based clustering perspectives, in which “true” states are defined relative to study aims, temporal resolution, and covariate design rather than as unique, data-determined entities (Hennig, 2015). Finally, we determine the effective number of states via simple penalties on the transition structure and on state occupancy/labelling (minimum prevalence, persistence bounds, and refitting when empty states are detected), in contrast to approaches that rely on shrinkage or regularisation of state-specific coefficients (Dupont et al., 2025).

In summary, the suggested framework advances behavioural state inference from animal tracking data by integrating ecological theory, formal equivalence with movement models, and practical aspects such as control sampling and state persistence. Its combination of flexibility, interpretability, and statistical rigour makes it a powerful and generalisable tool for movement ecology.

## Acknowledgements

This work was supported by the Sentinel North program of Laval University, funded by the Canada First Research Excellence Fund. D.F. was also supported by the Natural Sciences and Engineering Research Council of Canada (NSERC). We acknowledge the technical support and computing infrastructure provided by Calcul Québec and Compute Canada. We also thank Louis-Paul Rivest, Natasha J. Klappstein, and Théo Michelot for insightful discussions.

## Conflict of interest statement

All authors declare that they have no conflicts of interest.

## Author contributions

I.B. and D.F. conceived the project idea. I.B. and A.N. designed the framework. I.B. developed the models and simulations, conducted the analyses. D.F. secured funding. I.B. led the writing of the manuscript. All authors provided critical feedback on the manuscript and approved the final version for publication.

## Data availability statement

The dataset for the plains zebra (Equus quagga) was obtained from Michelot et al. (2019) and Klappstein et al. (2023)), and is available at: (DOI: 10.5281/zenodo.7872602). The source code of the simulation and the application is available [here](https://datadryad.org/share/LINK_NOT_FOR_PUBLICATION/ysKR8aQdc7O-41xiQj8QVfQvNFSZUPsyxvx6xK-sLsA).

## Supporting information

**S1. The general structure of the framework for** *K*_*max*_ = 4.

**S2. HMM-SSF model for plain zebra**.

## Notes

### Competing Interest Statement

The authors have declared no competing interest.

